# Fetal network controllability co-develops with synaptic development and synchronizes with maternal network controllability during pregnancy

**DOI:** 10.1101/2025.09.03.674103

**Authors:** Huili Sun, Takuya Toyonaga, Saloni Mehta, Huaigui Liu, Chris Camp, Samantha Rossano, Krista Fowles, Sean Martins, Marisa Spann, Stephanie Groman, Richard E. Carson, Dustin Scheinost

**Affiliations:** Department of Biomedical Engineering, Yale University, New Haven, CT; Department of Radiology & Biomedical Imaging, Yale School of Medicine, New Haven, CT; Department of Radiology, Tianjin Key Lab of Functional Imaging & Tianjin Institute of Radiology, Tianjin Medical University General Hospital, Tianjin, China; Interdepartmental Neuroscience Program, Yale School of Medicine, New Haven, CT; Life Molecular Imaging, New York, NY; Department of Psychiatry, Vagelos College of Physicians and Surgeons, Columbia University, New York, NY; Neuroscience Institute, University of Chicago, Chicago, IL; Department of Psychiatry, Yale School of Medicine, New Haven, CT; Department of Anesthesia and Critical Care, University of Chicago, Chicago, IL; Wu Tsai Institute, Yale University, New Haven, CT; Yale Biomedical Imaging Institute, Yale School of Medicine, New Haven, CT; Department of Statistics and Data Science, Yale University, New Haven, CT; Child Study Center, Yale School of Medicine, New Haven, CT

## Abstract

White matter undergoes rapid changes during the fetal period that are foundational for future cognitive functions. However, how these changes contribute to the brain’s capacity to support its dynamic activities—its *controllability*—remains largely unknown. Here, we apply network control theory (NCT) to investigate the developmental trajectory of controllability from the second trimester through the first postnatal month. We analyzed structural connectivity data from fetuses and infants as part of the developing Human Connectome Project. We identified a robust, nonlinear U-shaped developmental curve of whole-brain controllability across the perinatal period, with a minimum at approximately 35 weeks of gestation. Preterm birth disrupted these trajectories, leading to greater controllability and earlier minimums compared to age-matched fetuses. Using gene expression microarray data from 18 fetal post-mortem brains, we identified genes implicated in synaptic functions that co-develop with changes in controllability during the fetal period. We then used positron emission tomography in seven pregnant rhesus macaques to quantify changes in fetal synaptic density. Increased synaptic density in non-human primates (NHPs) co-occurred with periods of reduced controllability in humans. Finally, using longitudinal scans of a pregnant woman, we mapped the trajectory of changes in maternal controllability during pregnancy. This trajectory exhibited a U-shaped pattern that inversely correlated with the fetal trajectory, reaching a maximum around 36 weeks. Together, fetal controllability follows a nonlinear trajectory that co-develops with synaptic functions and synchronizes with maternal changes in controllability during pregnancy.

## Main

The brain undergoes tremendous functional and structural growth during the perinatal period. At the cellular level, synaptogenesis, myelination, and other processes begin prenatally, laying the foundation for the formation of complex functional networks at birth^1^. For example, major white-matter tracts are present in the third trimester^2^, leading to the establishment of highly connected hub regions and an adult-like network organization by birth^3^. Functional networks develop in the third trimester, initially forming homotopic connectivity and subsequently establishing longer anteroposterior connections^4^. At birth, many canonical resting-state networks are observable^5^. While structure and function are often studied independently during the perinatal period, white matter constrains functional activity, dynamics, and network structure. However, the applications of formal computation theory to understand how structural connectivity supports functional dynamics in fetuses and infants are nascent.

Network control theory (NCT) provides a formal framework for understanding how functional dynamics emerge from the structural connectome^6^. Controllability, an energy-based measure, quantifies the ability of gray matter regions to drive transitions between brain states^7^. In other words, controllability is a graph theory approach that explains how structural network topology informs and constrains dynamics. The maturation of controllability is an important milestone at several developmental stages. For example, during adolescence, structural connectivity and controllability mature, reducing the theoretical energy required for transitioning between activation states that are essential for adult-level executive function^8^. Similarly, controllability develops rapidly in the perinatal period, with elements of controllability observed in the third trimester and firmly established after term birth^9^. Nevertheless, this initial work in infants had several limitations. Most importantly, inferences of controllability before term birth were based on preterm infants, rather than fetuses. Preterm infants exhibited altered controllability and trajectories at term equivalent age and do not represent normative development. Preterm infants also show a wide range of brain differences compared to age-matched fetuses^10^. Finally, all preterm infants were scanned at 28 weeks of gestation or older, missing earlier development in the second trimester. Overall, the developmental trajectories of controllability in the perinatal period remain incomplete without normative fetal data.

Here, we created a normative growth curve of whole-brain controllability through the perinatal period in fetuses and infants. We then utilized this normative model to examine the associations between controllability, preterm birth, synaptic density, and maternal brain changes. Overall, we leveraged data from 217 fetuses, 448 term infants, and 194 preterm infants from the developing Human Connectome Project (dHCP)^11^, gene expression microarray data from 18 fetal post-mortem brains from the BrainSpan Atlas of the Developing Human Brain^12^, positron emission tomography (PET) with synaptic vesicle glycoprotein 2A (SV2A) radioligands in seven pregnant rhesus macaques based on previously developed method^13^, and one densely scanned pregnant woman from Pritschet et al.^14^ We find that controllability during the perinatal period follows a U-shaped trajectory, is altered by preterm birth (possibly in an experience-driven manner), associates with changes in synaptic gene expression, and inversely correlates with controllability in the maternal brain. Our results provide a template for and spur on future studies with multimodal, cross-species design and analysis, ranging from macroscale brain networks to molecular targets and gene expression, to generate a comprehensive understanding of brain development in the context of pregnancy.

## Results

We quantified controllability of the structural connectome for 217 fetuses (98 females, 119 males), 448 term infants (209 females, 239 males), and 194 preterm infants (86 females, 108 males) from the developing Human Connectome Project (dHCP; Table 1). Fetuses were scanned between 20.96 and 38.29 gestational weeks. Preterm infants were born between 23.00 and 36.86 weeks of gestation and scanned between 26.71 and 45.14 weeks. Term infants were born between 37.00 and 42.29 weeks of gestation and scanned between 37.43 and 44.71 weeks. We used DSI-studio (http://dsi-studio.labsolver.org/) to process the diffusion-weighted imaging data using generalized q-sampling imaging and create structural connectomes using the mean quantitative anisotropy value for an infant-specific atlas of 90 nodes^15^ (Fig. 1). Controllability was operationalized as average controllability, or the ability to drive the brain toward nearby brain states, and was calculated for each region in the connectome for each participant. Whole-brain controllability was defined as the mean across all regions in a single participant. Network-level controllability was defined as the mean across all regions in one of eight canonical networks^16^ (e.g., the default mode network) in a single participant. These data were used to establish normative trajectories of controllability throughout the perinatal period. These normative curves were used to investigate associations among gene expression in 18 fetal post-mortem brains and changes in controllability in the maternal brain in a single, densely scanned pregnant woman (Fig. 1).

**Fig. 1.**
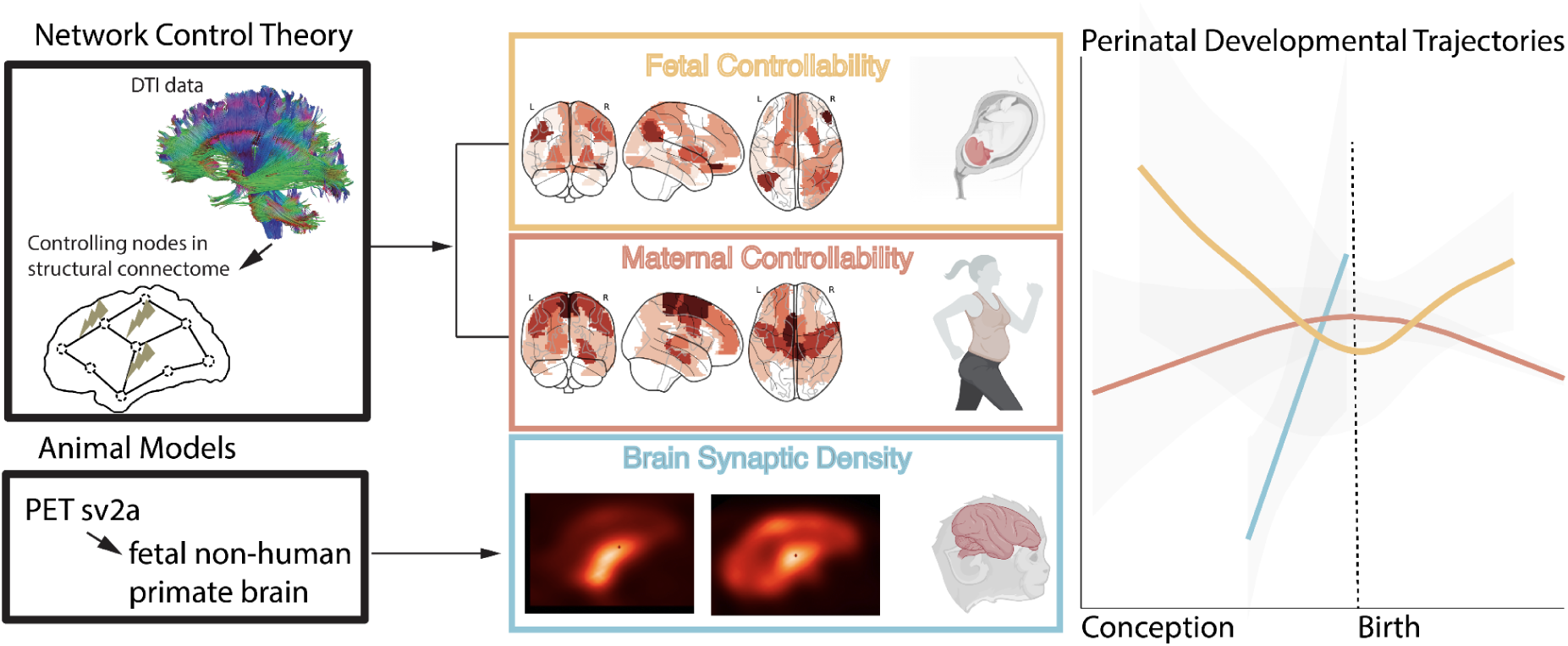
Study overview. Using network control theory, we quantified controllability in over 800 fetuses and infants and created normative curves from 20 to 45 weeks. A schematic of this curve is shown on the right in yellow. Using PET with SV2A radioligands in seven pregnant rhesus macaques, we mapped changes in synaptic density between 109 and 152 days of gestation. A schematic of this curve, projected onto the human gestational timeline, is shown in blue. Finally, we mapped changes in controllability during pregnancy using data from single women. A schematic of this curve is shown in red.

### Controllability exhibits a U-shaped developmental trajectory in the perinatal period

We first characterized changes in controllability across the perinatal period, from 20.86 to 45.14 weeks of gestation. We regressed sex, head motion, and network strength to normalize the fetal and infant data. Developmental trajectories were created using Generalized Additive Models for Location, Scale, and Shape (GAMLSS). Models revealed a strong non-linear association between gestational age and whole-brain controllability (F=77.01, p<2e-16, Deviance explained 38.6%, adj R-sq=0.382; Fig. 2). Whole-brain controllability decreased during the second and early third trimester, reached a minimum at 35.24 weeks, and increased through infancy. Next, we investigated the development trajectories for the controllability of canonical networks. Most networks showed similar non-linear trends. Two networks showed contrasting patterns. The ventral attention network showed no significant changes, whereas the limbic network exhibited an opposite, nonlinear trend (Fig. 2; Table S1).

**Fig. 2.**
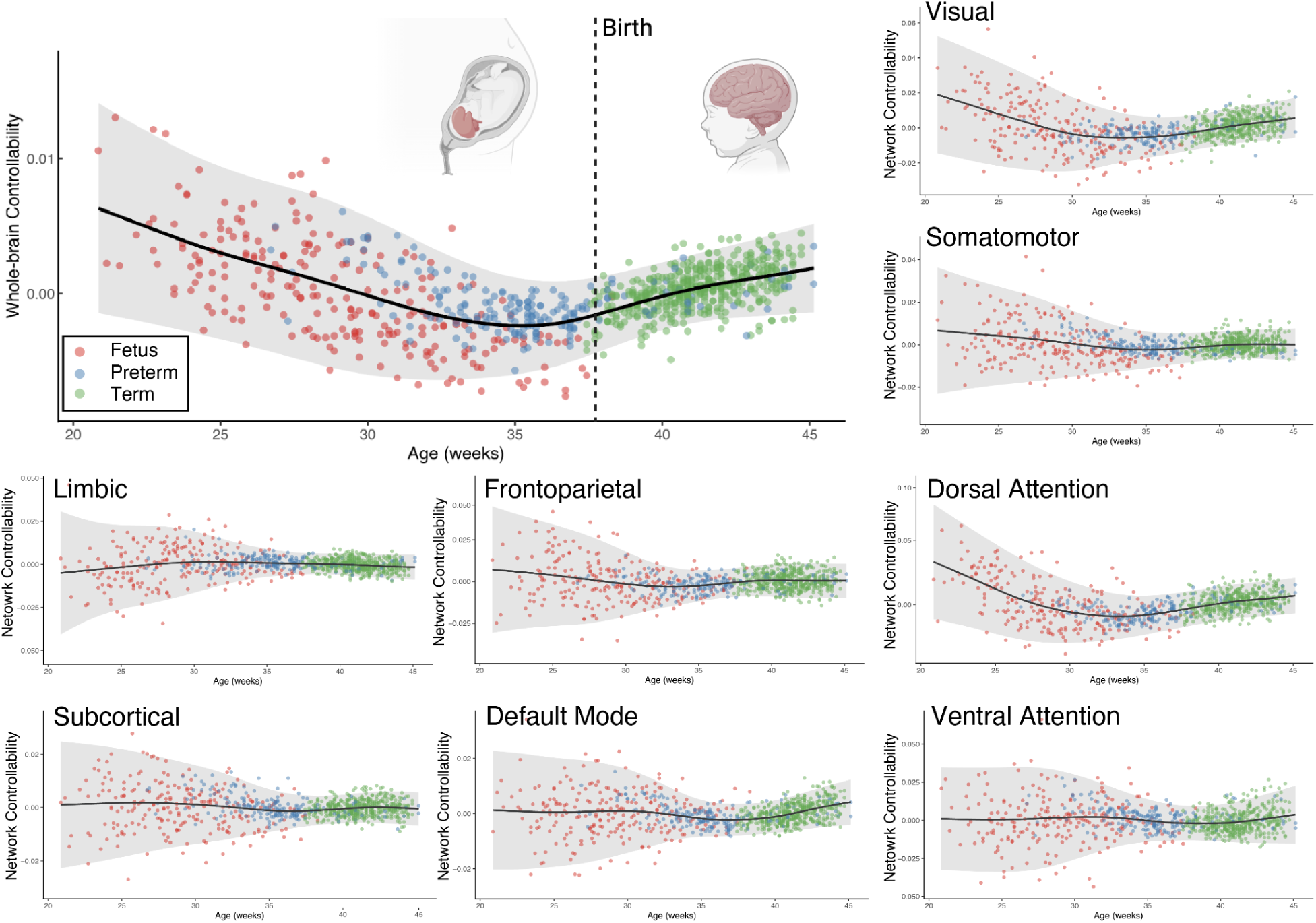
Controllability trajectories during the perinatal period. Generalized Additive Models for Location, Scale, and Shape (GAMLSS) were used to model the development of controllability at the whole-brain and network levels. A non-linear association was observed between gestational age and whole-brain controllability between 20 and 45 weeks of gestation. The curve reached a minimum at 35 weeks. Generally, the controllability of individual networks exhibited similar trends.

### Preterm infants exhibit greater controllability than age-matched fetuses

Next, we examined how preterm birth impacts controllability in the third trimester by comparing fetuses to preterm infants at matched gestational age on a week-by-week basis (Fig. 3a, Table S2). Across the third trimester (29 to 37 weeks), preterm infants exhibited significantly greater (p<0.05 corrected) whole-brain controllability. Preterm infants also reached their minimum sooner at 35.4 weeks compared to fetuses, who reached their minimum at 37.4 weeks. When compared at the regional level, greater controllability was generally observed across the whole brain in preterm infants (Fig. 3b). Overall, preterm infants demonstrate controllability curves that are both greater and more rapid in reaching their inflection point. Additionally, these results highlight how only using preterm infants to study brain development over the third trimester can bias and paint an incomplete picture of developmental trajectories.

**Fig. 3.**
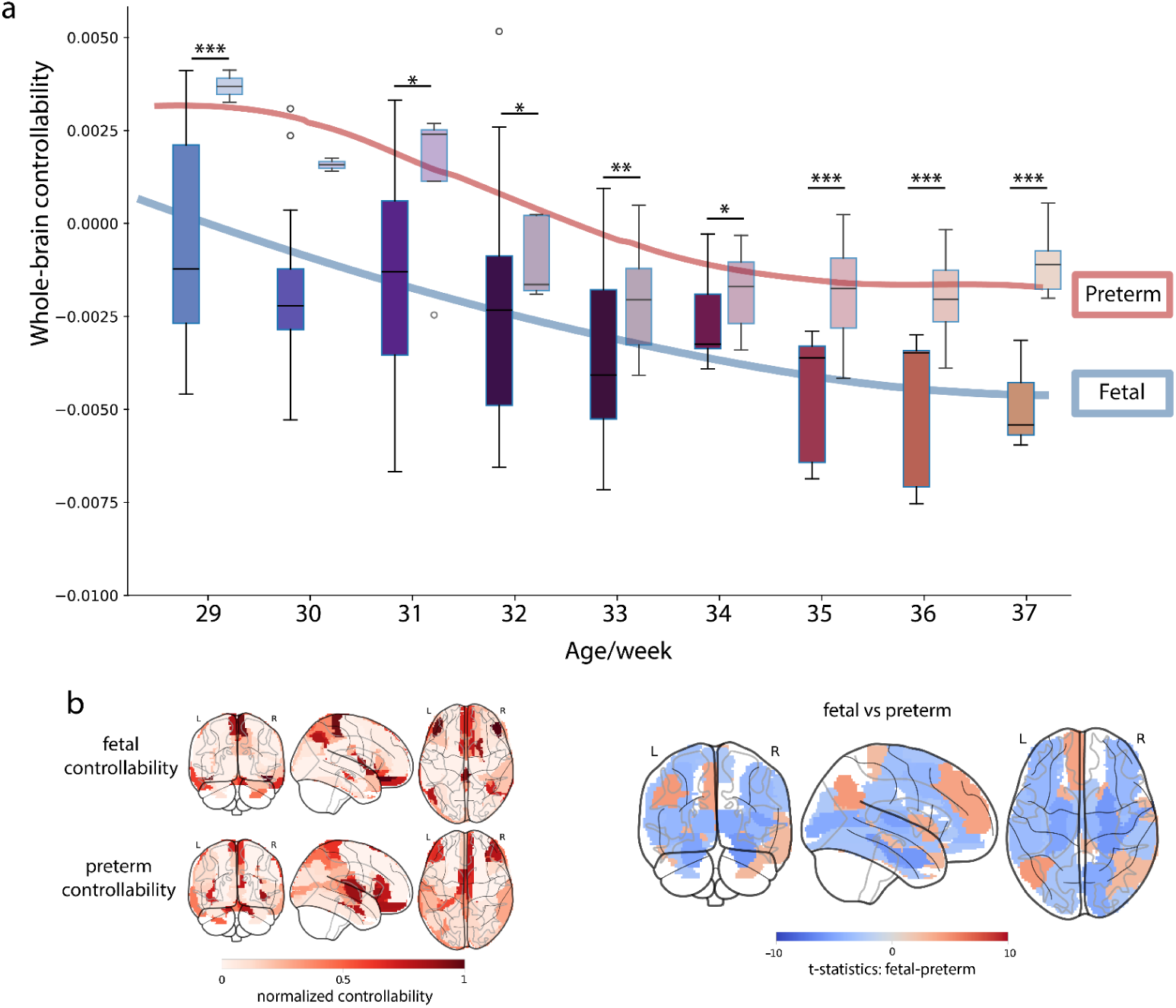
Differences in controllability between fetuses and preterm infants at matched gestational ages. (a) Group differences between preterm infants (left box) and fetuses (right box) at matched gestational ages on a week-by-week basis. Group differences (p<0.05, FDR-corrected) were observed during the third trimester. (b) Brain maps for normalized controllability in fetal and preterm infants. (c) Regional differences in controllability between fetuses and preterm infants between 29 and 37 gestational weeks.

### Reversed developmental patterns in controllability are observed in fetuses and infants

In general, whole-brain controllability reached a minimum around full-term birth. We investigate how developmental patterns differed between decreases in controllability during the fetal period and increases in controllability during the infant period using predictive modeling. We first trained a linear model to predict gestational age in fetuses using regional controllability and 10-fold cross-validation (Fig. 4). This model demonstrated significant predictions (r=0.80, p=8.02e-54, MAE=1.87 weeks). Similarly, we trained a linear model to predict postmenstrual age in term and preterm infants, yielding significant predictions (r=0.87, p=5.68e-198, MAE=1.39 weeks). The brain masks used in the predictive models are shown in Fig. S1. Next, we generalized the models between fetuses and infants. As different slopes are observed for fetuses and infants (Fig. 2), the sign on all model weights was multiplied by negative one. These models showed significant generalization (fetal to infant: r=0.59, p=4.78e-61, MAE=8.31 weeks; infant to fetal: r=0.61, p=8.56e-25, MAE=5.38 weeks). These results suggest that fetuses and infants exhibit a reversed pattern of regional controllability during maturation. In other words, the regions that decrease in controllability during the fetal period are the same regions that increase in controllability during infancy.

**Fig. 4.**
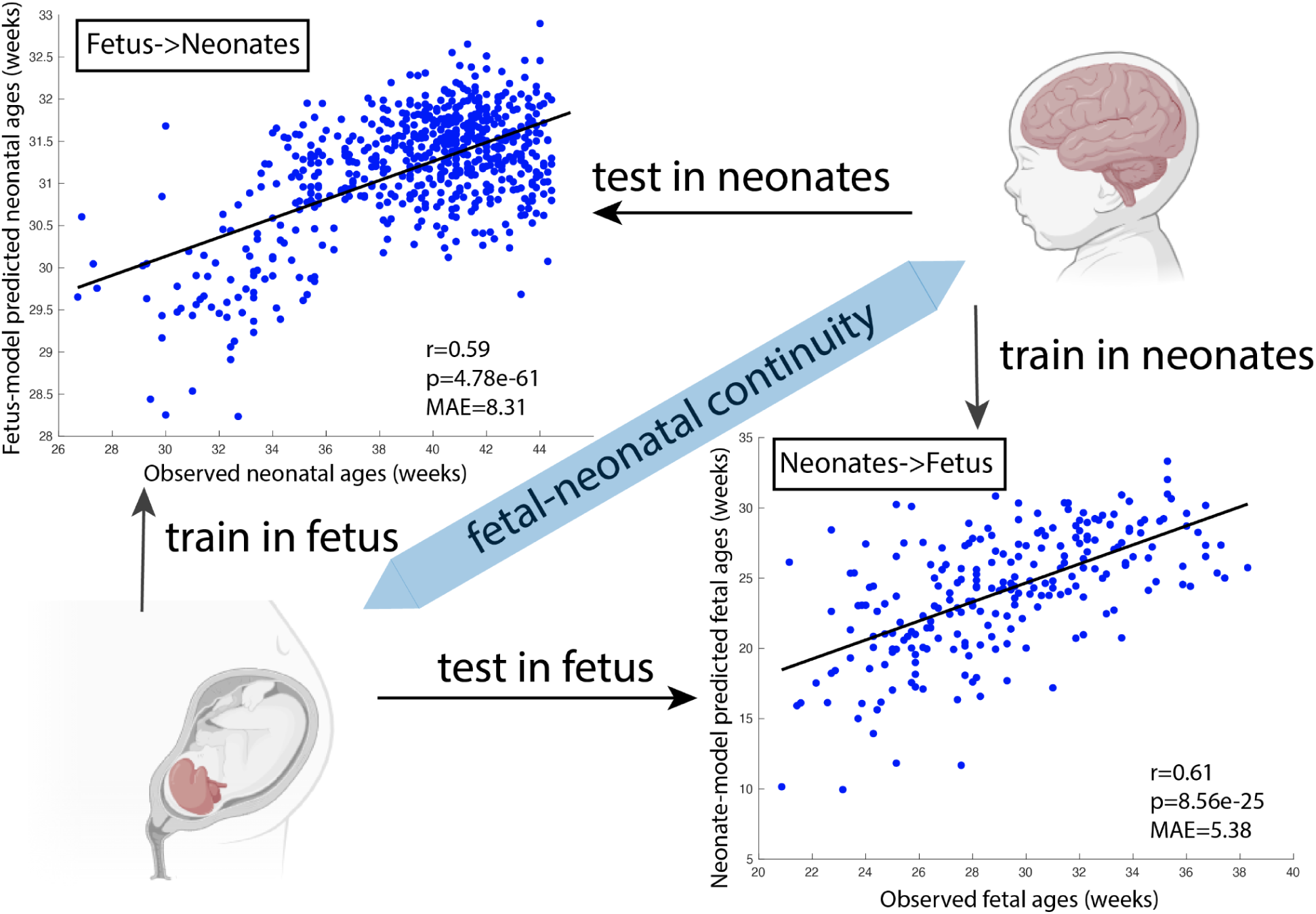
Cross-prediction of ages between fetal and infant groups. Two predictive models of age were trained independently on fetal and infant controllability data. Both the fetal and infant models accurately predicted age. Next, we reserved the signs on the models’ weights to account for the different slopes of controllability during the fetal and infant periods (Fig. 2) and generalized the models across developmental stages. The infant model significantly predicted fetal gestational age (r=0.59, p=4.78e-61; upper-left scatter plot). The fetal model significantly predicted infant postmenstrual age (r=0.61, p=8.56e-25; lower-right scatter plot).

### Synaptic development may underlie the changes in controllability during the fetal period

We next investigated the possible biological mechanisms underlying changes in fetal controllability. To determine potential mechanisms that co-develop with controllability, we screened for genes with expression trajectories similar to those of controllability, using data from the transcriptional landscape of the fetal brain^12^ (https://www.brainspan.org/). In the module C22, which was previously detected as corroborated with cortical development^12^, we found several genes (i.e., *TMEM59L*, *CAMK1*, *PINK1*, *PRRT3*, and *CKMT1A*) whose expression changes over the fetal period significantly correlated with the changes of controllability after Bonferroni-Holm correction for multiple comparisons (*TMEM59L*: r=-0.97, p=8.87e-4, *CAMK1*: r=-0.96, p=0.0029, *PINK1*: r=-0.93, p=0.017, *PRRT3*: r=-0.92, p=0.038 and *CKMT1A*: r=-0.94, p=0.0097; Fig. 5a). Permutation testing (1,000 iterations with shuffled gene expression trajectories) yielded no significant correlations with fetal controllability (Fig. S2). A majority of these genes are enriched in the brain, with their expression clustered in synaptic function (Table S3).

**Fig. 5.**
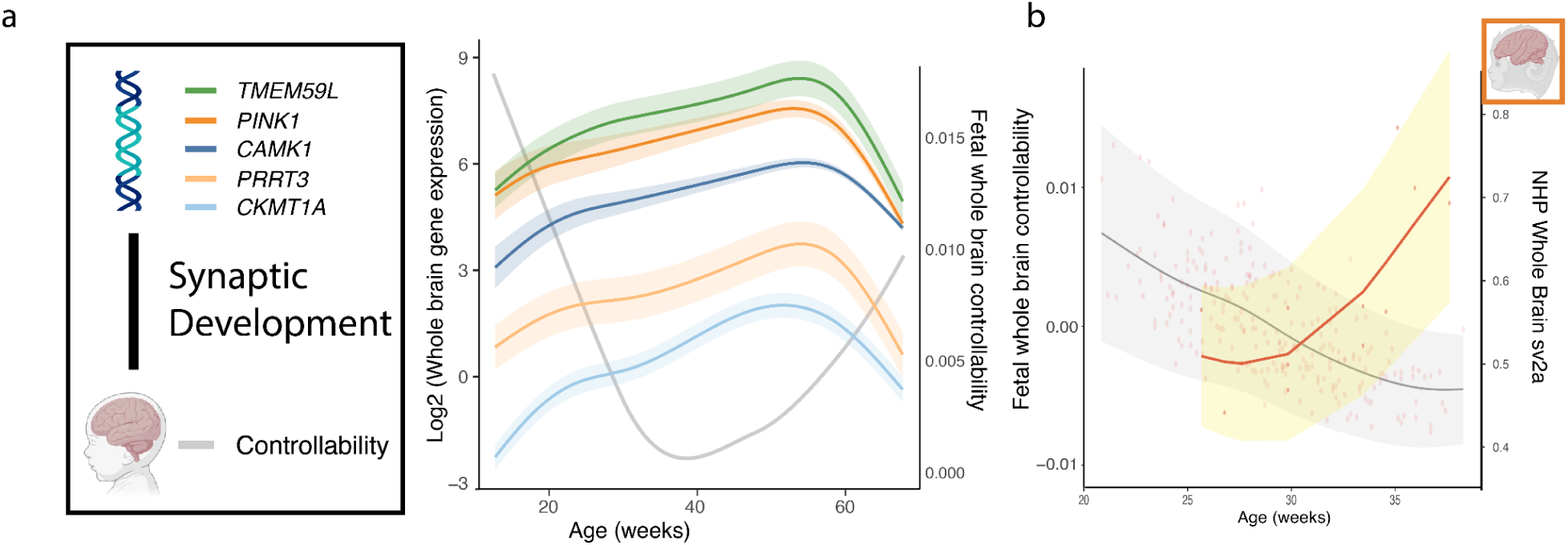
Associations between synaptic development and brain controllability. (a) Gene expression trajectories from the fetal brain transcriptome (BrainSpan) were screened against the developmental curve of controllability, revealing significant correlations for genes enriched in synaptic function (e.g., *TMEM59L*, *PINK1*, *CAMK1*, *PRRT3*, and *CKMT1A*). (b) SV2A concentration in NHP fetal brains increased over the 3rd trimester, suggesting rapid synaptogenesis.

Next, we investigated in vivo measures of synaptic density in the brain using positron emission tomography (PET) with [^11^C]UCB-J, a SV2A radioligand, in the fetuses of seven pregnant rhesus macaques. SV2A is a glycoprotein located on the vesicles in presynaptic terminals and is an indirect estimate of synaptic density^17^. Each was between 109 and 152 days of gestation (term pregnancy: 165 ± 10 days). Whole-brain distribution volume ratio (DVR), as a measurement of SV2A density, increased across the third trimester (β=0.042 DVR/week, p=1.24e-4, SE=0.0070, 95% CI=[0.0268, 0.0580]; Fig. 5b). Regional associations were similar (Table S4). In contrast, whole-brain DVR in the maternal brain did not change over pregnancy (Table S5; Fig. S3).

### Maternal controllability trajectory inversely correlates with fetal-infant trajectories

As pregnancy also witnesses profound changes in the maternal brain^14,18^, we investigated the trajectory of maternal controllability across pregnancy using precision neuroimaging data from Pritschet et al.14 These data consisted of 20 scanning sessions from a 38-year-old woman spanning pregnancy and the early postpartum period. Models exhibited a strong non-linear association between the week of pregnancy and whole-brain controllability (F=4.38, p=0.017, Deviance explained 47.30%, adj R-sq=0.40; Figure 6). Whole-brain controllability increased from the start of pregnancy, reached a maximum at 36.63 weeks, and then decreased through the postpartum period. These curves broadly mimic the trajectories of other structural neuroimaging features^14^. At the network-level, the somatomotor, dorsal attention, frontoparietal, and subcortical networks exhibited significant trajectories with a similar shape to that of the whole-brain (Table S6; Fig. S4).

**Fig. 6.**
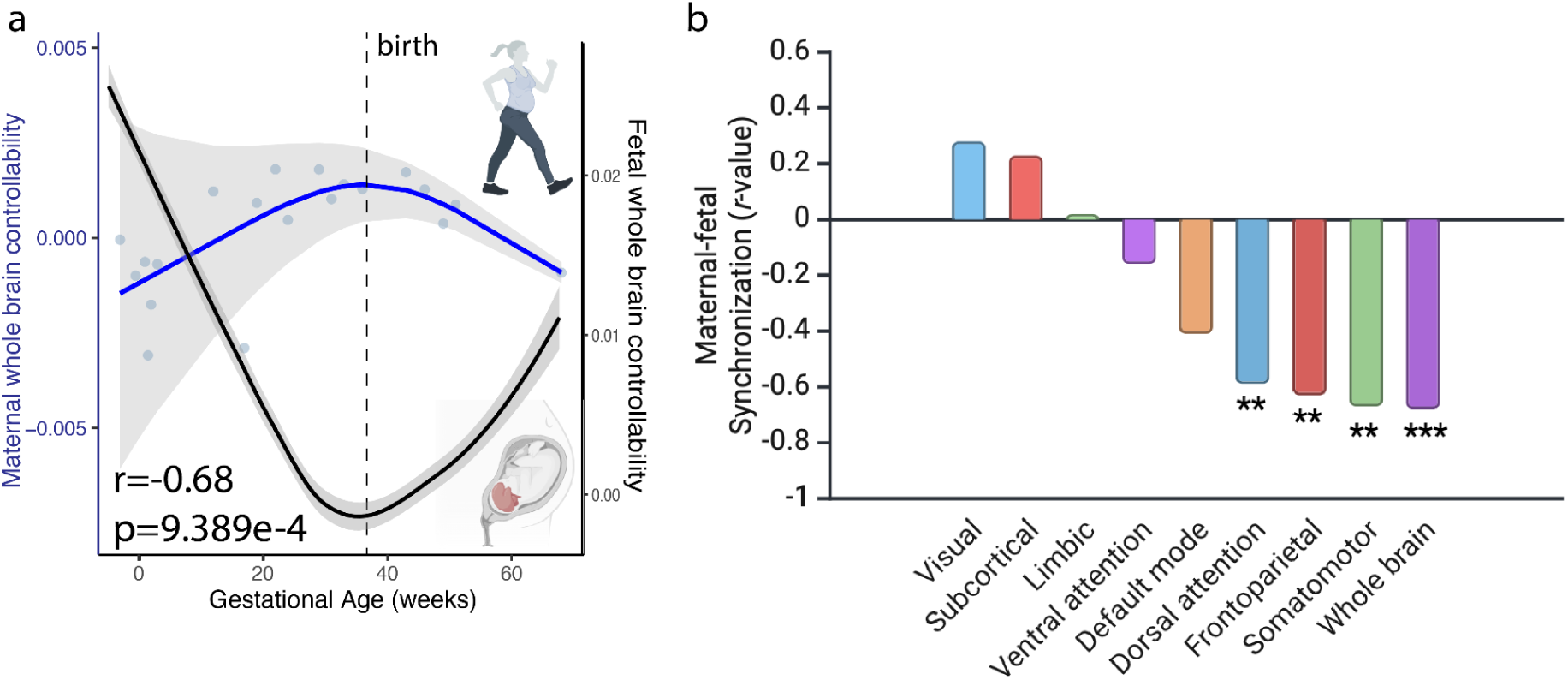
Maternal-fetal synchronization in brain controllability changes. (a) On the whole-brain level, controllability in the maternal brain changed in an opposite trend compared to that of the fetal brain. (b) Significant anti-correlations between maternal and fetal brain controllability were also observed in the dorsal attention, frontoparietal, and somatomotor networks.

Next, we compare the maternal controllability trajectories (Fig. 6a) to the fetal-infant ones (Figure 2). Using the fetal-infant GAMLSS model, we generated normative fetal and infant controllability values at the same imaging time points as the maternal imaging data. Next, we correlated these predicted fetal values with the measured maternal controllability on both whole-brain and network levels. At the whole-brain level, we observed an inverse correlation between the two trajectories (r=-0.68, p=9.39e-4). Notably, both curves achieved their extremum at similar times (maternal maximum at 36.63 gestational weeks; fetal minimum at 35.24 gestational weeks). At the network level, the somatomotor, dorsal attention, and frontoparietal networks exhibited similar, significant (p<0.05, corrected) associations (Fig. 6b).

## Discussion

Applying network control theory to fetal and infant neuroimaging data, we charted the development of controllability during the perinatal period. Controllability decreased during the second trimester, reaching the minimum shortly before full-term birth, and then increased during early infancy. Preterm birth disrupts this trajectory, leading to greater controllability and earlier minimums compared to age-matched fetuses. Fetuses and infants exhibit a reversed pattern of regional controllability during maturation, in which the regions that decrease in controllability during the fetal period are the same ones that increase in controllability during infancy. These changes in brain controllability were potentially driven by underlying synaptic development, as suggested by changes in NHP PET data and significant associations with synaptic gene expression. Additionally, controllability in the maternal brain followed an opposite trend to that in the fetal brain. Together, we comprehensively characterize the developmental trajectory of network controllability during the perinatal period. Moreover, associations between these trajectories and maternal brain changes during pregnancy, as well as molecular imaging in NHPs, provide a broader perspective on these developmental patterns.

The U-shaped trajectory of controllability reaches a minimum around full-term birth. Dendritic arborization and synaptogenesis accelerate in the third trimester, rapidly increasing the complexity of early neural circuits^10^. This growing complexity may underlie the decreasing controllability during the fetal period, as more complex circuits are less efficient at controlling functional dynamics. Our NHP results support with this interpretation. This inverse correlation aligns with adolescent literature, where controllability increases during adolescence^19^ and synaptic pruning is pronounced, with some regions losing 50% of their synaptic connections^20^. Though no studies have directly linked these effects during adolescence. Our transcriptional results provide further evidence for synaptic development as a potential mechanism underlying changes in controllability^21,22^.

The increase in controllability around birth may be attributed to several factors, including myelination. While myelination begins prenatally, the majority of white matter remains unmyelinated in fetuses and increases fivefold by full-term birth. As myelination increases the speed and efficiency of neuronal communication, the ability to control functional dynamics should also increase. Further, this increased myelination may moderate the association between synaptic development and controllability, such that it is reduced with sufficient myelination. In addition, the differences between the in-utero and ex-utero environment likely contribute to this U-shape. Newborns experience a rapid increase in sensory stimuli after birth. These experiences help refine neural functions and establish cognitive circuits^23^. They likely also facilitated the growth in controllability after birth^9^. This hypothesis is supported by our preterm results, which showed an earlier minimum and greater controllability compared to age-matched fetuses. Preterm infants are exposed to ex-utero stimuli earlier and thus have greater controllability until full-term birth^10,24^. The greater controllability, in turn, may lead preterm infants to exhibit various skills earlier than fetuses.

Preterm birth exerts a significant impact on controllability compared to healthy fetuses over a wide range of gestational ages. Aligned with this finding, alterations in brain structure are widely reported in preterm birth compared to fetal brains at the same gestational age^25–27^. Fetal brain structure changes consistently before 30 weeks of gestation, where the fractional anisotropy constantly decreases and mean diffusivity increases in most neural pathways. These trends slow down and even reverse before birth^28^. Preterm birth in the late third trimester may contribute to the onset of the reversal process faster than usual, leading to a more significant effect on mid- and later-preterm infants.

A mother and her fetus form a dyad that interacts dynamically and bidirectionally. This dyadic synchrony has been studied using heart rate and motor activity^25,29^, particularly in response to external stimuli, such as the mother’s voice^30^. Indeed, hyperscanning of parents and older infants also reveals neural synchrony during social interactions^31^. In alignment, we observed synchrony in changes in controllability between the mother and the fetus, with both showing extrema at similar times. Notably, an inverse association was observed, suggesting conservation across the dyad. For example, the dyad may not be able to support simultaneous increases in controllability across two brains. Although neuroimaging studies in fetuses and pregnant women are emerging^14,32^, studies that scan both simultaneously or in close temporal proximity do not exist. Such studies are logical next steps to understand how dynamic brain changes in the maternal-fetal dyad correlate and influence each other.

Molecular neuroimaging via PET is rare in early life. PET uses radioactivity isotopes to trace specific molecular targets and, thus, is unsuited for research in pregnant women and infants. NHP models represent a novel approach to linking molecular neuroimaging with more conventional MRI measures currently collected in humans. In the future, PET and MRI can be collected in NHP fetuses (potentially simultaneously with PET-MRI), allowing the same MRI measures to be collected in both species. Additionally, the experimental paradigms available in NHP are a natural complement to observational studies typically used with human fetuses and infants. Combining these studies enables the investigation of potential cellular and molecular mechanisms of many well-known prenatal exposures^33^ in cross-species translational studies.

Our work has several strengths, including multimodal neuroimaging data from fetuses, a large cohort of fetal and infant data, state-of-the-art computational modeling (GAMLSS, NCT, predictive modeling), cross-species translation, gene expression, and a combination of maternal and fetal neuroimaging data. However, limitations exist. First, differences in fetal and infant imaging protocols may influence results, even though these data were collected on the same scanner. For example, the dHCP study utilized a cardiac coil for fetal scans^34^ and a dedicated head coil for infants^11^. We regressed sex, head motion, and network strength to minimize these effects. Additionally, although preterm infants differ from normative development, they exhibited broadly similar trajectories (i.e., decreasing controllability until the late third trimester and increasing thereafter). They were scanned with the same protocol as the term infants. Together, these suggest that the observed trajectories are not due to systematic scanning differences. Second, we used a conventional PET/CT (Siemens Biograph) for synaptic density measurements. It and similar scanners have inherent limits for spatial resolution and sensitivity, making it suboptimal for capturing fine-grained SV2A changes in small brains, such as fetal NHPs. Next-generation imaging approaches, such as the NeuroExplorer, could provide more definitive insights through its enhanced spatial resolution and sensitivity^35,36^. Third, all data were from secondary data analyses and not designed for the investigations at hand. As a result, comparisons across datasets were based on normative values from our GAMLSS models, which limits the strengths of the conclusions. For example, our NHP sample lacked MRI data. Collecting fetal MRI data from the NHP sample would allow direct comparisons of controllability and synaptic density within the same animal. It could also facilitate more precise gestational mapping, aligning controllability trajectories in NHPs and humans. Similarly, fetal and maternal brain data were not from the same dyad. Nevertheless, both of these datasets are one of a kind. Our results highlight the potential of secondary data analysis to inspire new data collection in innovative study designs, thereby addressing the limitations of our work.

In conclusion, we characterized the developmental trajectory of controllability through the perinatal period. These trajectories co-occur with synaptic-related gene expressions and synaptic density changes witnessed in NHPs, and our results highlight the synchrony of brain changes in the maternal-fetal dyad. Insights into normative brain development might open the door to early identification and intervention methods to improve developmental outcomes.

## Methods

### MRI scans from dHCP

Fetal and neonatal neuroimaging data were obtained from the Developing Human Connectome Project^37^ (dHCP, http://www.developingconnectome.org/), a cross-sectional open-source study of infant brain development. The National Research Ethics Service, West London committee, approved the study. Participating families provided written consent before imaging. Our sample included 807 subjects: 217 fetuses (98 females, 119 males), 448 term infants (209 females, 239 males), and 194 preterm infants (86 females, 108 males) from the second data release of dHCP. Fetuses were scanned between 20.96 and 38.29 gestational weeks. Preterm infants were born between 23.00 and 36.86 weeks of gestation and scanned between 26.71 and 45.14 weeks. Term infants were born between 37.00 and 42.29 weeks of gestation and scanned between 37.43 and 44.71 weeks.

All images were collected in the Evelina Newborn Imaging Center, St Thomas’ Hospital, London, UK. MRI data were acquired using a Philips Achieva 3 T scanner (Philips Medical Systems, Best, The Netherlands) for both fetal and neonatal participants^34^. Fetal participants were scanned using a 32-channel cardiac coil. Sequences were extensively tested and optimized with safety constraints relevant to fetal scanning, which are paramount, namely, restricting SAR (<50%), PNS (<60%), and acoustic noise (<110 dB). 2D snapshot multi-slice acquisitions are used for all modalities to freeze maternal-induced motion and spontaneous fetal movements. Structural T2w data is acquired from 6 uniquely oriented stacks centred to the fetal brain using a zoomed multiband (MB) single-shot TSE sequence (TE = 250 ms, TR = 2265 ms, resolution = 1.1 x 1.1 x 2.2 mm^3^ with a 1.1 mm gap) with a MB tip-back preparation pulse for increased SNR efficiency^38^. Each anatomical dataset is reconstructed into an isotropic 3D volume using slice-to-volume reconstruction (SVR), with automatic rejection of corrupted data^39^. A neuroradiologist reported that the brains of all fetuses showed appropriate appearances on the T2-weighted anatomical scan for their GA, with no acquired lesions or congenital malformations of clinical significance. Diffusion data consisting of 141 volumes (15 b=0 s/mm^2^, 46 b=400 s/mm^2^, and 80 b=1000 s/mm^2^, designed using a data-driven method) is acquired using a spin- and field-echo (SAFE) sequence. dMRI suffered from time-varying EPI distortion due to B0 disturbance caused by maternal breathing. The inherent phase of the slice data, which combines spin-echo and field-echo phase information for dMRI, is used to estimate dynamic B0 field variation, thereby undistorting the images shot-by-shot, before motion correctio^40^. The SHARD pipeline was then used to correct motion and dMRI artifacts (e.g., eddy current, Gibbs ringing, and susceptibility artifact^41^.

A dHCP-customized neonatal imaging system, including a 32-channel receive neonatal head coil (Rapid Biomedical GmbH, Rimpar, DE) for infant subjects. Infants were scanned during unsedated sleep after feeding and immobilization in a vacuum-evacuated bag, with hearing protection and physiological monitoring (including pulse oximetry, body temperature, and electrocardiography data) applied during scanning. T2-weighted images were obtained using a Turbo spin echo sequence (TR = 12 s, TE = 156 ms, SENSE factor 2.11 (axial) and 2.54 (sagittal)) with overlapping slices (resolution = 0.8 × 0.8 × 1.6 mm^3^). T2w images were motion-corrected and super-resolved to a resolution of 0.8 × 0.8 × 0.8 mm^3^ ^42^. Diffusion-weighted imaging (DWI) was obtained in 300 directions (TR = 3.8 s, TE = 90 ms, SENSE factor 1.2, multiband factor 4, and resolution 1.5x1.5x3mm^3^ with 1.5 mm slice overlap) with b-values of 400 s/mm^2^, 1000 s/mm^2^, and 2600 s/mm^2^ spherically distributed in 64, 88, and 128 directions, respectively, using interleaved phase encoding.

Preprocessing of the data was performed to correct for noise^43^ and bias field inhomogeneities^44^. Registration of the images to the dHCP weekly T2 atlas^45^ was carried out using rigid transformation, and b-vectors were subsequently rotated accordingly. A quality check was conducted by the neighboring DWI correction (NDC)^46^, resulting in the exclusion of 12 fetal scans and 34 neonatal scans due to their low NDC values, as calculated by a median value-based outlier detector. The reconstruction of the diffusion data was performed in native space with generalized q-sampling imaging (GQI)^47^ with a diffusion sampling length ratio of 1.25. The tensor metrics were calculated and analyzed using the resource allocation (TG-CIS200026) at Extreme Science and Engineering Discovery Environment (XSEDE) resource^48^. After reconstructing images with GQI, the whole-brain fiber tracking was conducted with DSI-studio (http://dsi-studio.labsolver.org/) with quantitative anisotropy (QA) as the termination threshold. QA values were computed in each voxel in their native space for every subject. The tracking parameters were set to an angular cutoff of 60 degrees, a step size of 1.0 mm, a minimum length of 30 mm, and a maximum length of 300 mm. The whole-brain fiber tracking process was performed with the FACT algorithm until 1,000,000 streamlines were reconstructed for each individual^49^. Here, we used a neonatal AAL-aligned brain parcellation with 90 nodes to construct the structural connectome for each infant^15^. T2-weighted images in native DWI space were used to provide information on region segmentation during the construction of connectomes. The structural connectome for each individual was then constructed with a connectivity threshold of 0.001. The pairwise connectivity strength was calculated as the average QA value of each fiber connecting the two end regions, resulting in a 90 × 90 structural connectome for each participant.

### MRI scans during pregnancy

26 MRI scans of the human brain during pregnancy were obtained from a 38-year-old healthy primiparous woman from 3 weeks before conception through 2 years postpartum (162 weeks) as reported by Pritschet et al.^14^. Here, we focused on the 20 scans obtained during pregnancy and perinatal period, aligned with the human fetal and neonatal data. The participant experienced no pregnancy complications (for example, gestational diabetes and hypertension), delivered at full term via vaginal birth, nursed through 16 months postpartum, and had no history of neuropsychiatric diagnosis, endocrine disorders, prior head trauma, or history of smoking. The study was approved by the University of California, Irvine Human Subjects Committee. Written informed consent was collected.

MRI scanning sessions at the University of California, Santa Barbara and Irvine were conducted on 3T Prisma scanners equipped with 64-channel phased-array head/neck coil (of which 50 coils are used for axial brain imaging). High-resolution anatomical scans were acquired using a MPRAGE sequence (TR = 2,500 ms, TE = 2.31 ms, inversion time = 934 ms, flip angle = 7°, 0.8 mm thickness). A gradient echo field map was also collected (TR = 758 ms, TE1 = 4.92 ms, TE2 = 7.38 ms, flip angle = 60°). The Diffusion Spectrum Imaging (DSI) protocol sampled the entire brain with the following parameters: single phase, TR = 4,300 ms, echo time = 100.2 ms, 139 directions, b-max = 4,990, FoV = 259 × 259 mm, 78 slices, 1.7986 × 1.7986 × 1.8 mm voxel resolution. A custom foam headcase was used to provide extra padding around the head and neck, thereby minimizing head motion. Additionally, a custom-built sound-absorbing foam girdle was placed around the participant’s waist to attenuate sound near the fetus during second-trimester and third-trimester scanning.

We followed the same preprocessing pipeline, using QSIprep 0.16.1 with a Singularity container and the same parameter settings, to reconstruct the diffusion MRI scans. Mini-preprocessed scans were then reconstructed in the native space with generalized q-sampling imaging (GQI) with a diffusion sampling length ratio of 1.25. After reconstruction, whole-brain fiber tracking was conducted using DSI-studio with quantitative anisotropy (QA) as the termination threshold. QA values were computed in each voxel in their native space for every subject. The whole-brain fiber tracking process was performed with the FACT algorithm until 1,000,000 streamlines were reconstructed for each individual. Here, we used the AAL2 atlas to construct a structural connectome for each scan^50^. T1-weighted images in native DWI space were used to provide information on region segmentation during the construction of connectomes. The structural connectome for each scan was then constructed with a connectivity threshold of 0.001. The pairwise connectivity strength was calculated as the average QA value of each fiber connecting the two end regions, resulting in a 120 x 120 structural connectome for each scan.

### Network control theory

The brain can be considered a natural complex system. A way to understand the behavior of a complex system is to understand the mechanisms that control it, which involves driving the system to the desired state. Network control theory, aimed at addressing the problem of how to control a system consisting of nodes (brain regions) and edges (white matter tracts between brain regions), can illustrate the dynamic changes in the short term or the developmental trajectory in the long term that the human brain goes through, and distinguish the cognitive dysfunction in disorder groups.

A network system can be represented as a graph 𝐆 = (𝐕, 𝐄), where 𝐕 and 𝐄 are the vertex and edge sets, respectively. Let 𝑎_𝑖𝑗_be the weight associated with the edge (𝑖, 𝑗) in 𝐄 and define the weighted adjacency matrix of the graph 𝐆 as 𝐀 = [𝑎_𝑖𝑗_], where 𝑎^𝑖𝑗^ = 0 when 𝑎_𝑖𝑗_ ∉ 𝐄. Here, the individual structural connectome 𝐀 in ℝ^𝑛×𝑛^ is a symmetric and weighted adjacency matrix whose elements [𝑎_𝑖𝑗_] evaluate the strength of the white matter fiber connecting between region 𝑖 and region 𝑗 in the brain.

For the definition of the neural dynamic processes, we adopt the prior models that link brain structural networks to simplified brain dynamics. Although the evolution of brain activity occurs in a nonlinear manner, previous studies have demonstrated that simplified linear models can predict a significant portion of the variance in neural dynamics recorded by fMRI^51^. Therefore, we employ a simplified noise-free linear discrete-time and time-invariant network model: 𝐱(𝑡 + 1) = 𝐀𝐱(𝑡) + 𝐁_𝐾_(𝑡)𝐮_𝐾_(𝑡) , where 𝒙 denotes the brain state at a given time, and 𝐀 is the symmetric, undirected, and weighted adjacency matrix for the network. In our case, 𝐀 represents the structural connectome for each individual, whose element indicates the pairwise strength of the structural connection. The diagonal elements are set to zero. The input matrix 𝐁_𝐾_ identifies the control points 𝛫 in the brain, where 𝛫 = {𝑘_1_, ···, 𝑘_𝑚_} and 𝐁_𝐾_= [𝑒_𝑘1_··· 𝑒_𝑘𝑚_]. 𝑒_𝑖_ denotes the 𝑖th canonical vector of dimension 𝑁 , and input over time. 𝒖_𝐾_denotes the input control strategy over time.

We study the ability of a region to drive a dynamic system to a desired state, which is defined as controllability. Classic control theory provides that the controllability of the network from the set of network nodes 𝐾 is equivalent to the controllability Gramian 𝐖_𝐾_ being invertible, where 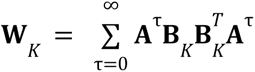. The input matrix 𝐁 reduces to a one-dimensional vector as we choose control nodes one at a time. Based on this network control theory framework, we examined average controllability, which is defined by the average input energy from a group of control nodes and overall potential target states. The average input energy is proportional to Trace (*W_K_*^−1^), the trace of the inverse of the controllability Gramian. Here, to maintain consistency with previous studies^19,52^, we use the Trace as the measure of *average controllability* to increase the accuracy of computation on small brain networks and preserve the information obtained from this measurement. As noted in the prior study, Trace(𝐖_𝐾_) encodes a well-defined metric for controllability, equivalent to the network response. 𝐻_2_ norm or the energy of the network’s impulse

### GAMLSS modeling

To outline the developmental trajectory of brain controllability during the perinatal period, we used a Generative Additive Model for Location, Scale, and Shape (GAMLSS; R package *gamlss*). Whole-brain and network brain controllability were modeled separately as a function of age using penalized B-splines for smoothing, with separate smoothing terms fitted for the location (mu), scale (sigma), skewness (nu), and kurtosis (tau) parameters under a normal distribution assumption. This approach enables age-dependent changes in both the mean trajectory and variability of controllability throughout gestation. Model selection was performed by comparing competing models, including models with reduced parameter sets and different distributional assumptions (normal vs. power exponential), using the Bayesian Information Criterion (BIC). Lower BIC values indicated better model fit. The fitted model was used to generate centile curves representing various percentiles of brain controllability across gestational age. Model predictions included z-scores for each observation, fitted values for all distributional parameters, and estimated slopes representing rates of change across the age range.

### Predictive modeling

To explore if the developmental pattern of controllability is shared between fetal and neonatal brains, we employed predictive modeling to predict age at scan for fetuses and infants. We trained the predictive model using brain controllability with validated MATLAB scripts. First, the fetal data were split into training and testing data using 10-fold cross-validation. Sex, brain volume, and head motion were regressed from each individual’s controllability. Features (i.e., regional controllability) significantly correlated (p<0.01) with age were selected in the training group. The selected features for each individual were summarized into a single number. Linear regression was then used to model this summary score and the ages in the training group. Finally, this model was applied to the testing group. Next, this model was generalized to the data from term and preterm infants. As the opposite pattern of controllability is observed before and after birth, the signs of the model were reversed to account for the change in trajectory. The procedure was repeated with 10-fold cross-validation on the infant data and then generalized to the fetal data. Prediction performance was evaluated using Pearson’s correlation (r) and mean absolute error (MAE).

### Nonhuman primate positron emission tomography (PET)

Seven gravid rhesus macaques were included. All animals, except one, were scanned at two gestational ages during the third trimester, once at approximately 120 days and once at approximately 145 days (term pregnancy: 165 ± 10 days). One rhesus macaque missed the second scan due to radiosynthesis failure. NHPs underwent PET scanning with [^11^C]UCB-J (160.8 ± 43.0 MBq) at the Yale PET Center (Yale School of Medicine, New Haven, CT). PET data were acquired for 0-60 min post-injection using the Siemens Biograph mCT (Siemens Medical Solutions, USA). Dams were sedated with ketamine/glycopyrrolate (7–10 mg/kg) and maintained under gas anesthesia (isoflurane; 0.75–2.5%) for the duration of the scan. PET images were reconstructed using time of flight (TOF) + point spread function (PSF) modeling and ordered subset expectation maximization (OSEM) algorithm with 21 subsets and 3 iterations. Fetuses were monitored sonographically across gestation, including immediately before each imaging session (around 120 and 145 days of gestational age). All procedures were carried out in strict accordance with the guidelines outlined in the Animal Welfare Act and the Guide for the Care and Use of Laboratory Animals.

In this study, both maternal and fetal SV2A PET uptake was investigated by using the maternal sentrum semiovale as a reference region. To determine regional SV2A PET uptake in the fetal brain, PET images were registered to the Neonate Rhesus Monkey template space^53^ for 2-week-old with a 12-degree-of-freedom transformation estimated using FSL^54^. The registration was guided by minimizing mutual information with an averaged fetal SV2A image acquired in the previous study that has been aligned to the same template space with 12 DoF^55^. For the maternal brain, a reference SV2A PET image was generated in template space using 10 previously acquired adult rhesus macaque [^11^C]UCB-J scans on the Focus 220 (Siemens). Transformations for those images were first estimated linearly between each PET and its corresponding individual MR, and then nonlinearly from each individual MR to the template MR^56^ . The 10 [^11^C]UCB-J PET images were then resliced into the template space, and the averaged image was created as the reference PET image. The maternal brain PET images in this study were registered to the reference PET image using a 12 DoF transformation. These estimated transformations were used to extract PET time-activity curves (TACs) in the following regions: frontal, parietal, occipital, temporal, insula, hippocampus, amygdala, caudate, putamen, and cerebellum, and centrum semiovale. Whole-brain SV2A was calculated as the average across all regions for each subject. Maternal centrum semiovale TAC was extracted and used as the reference region to estimate regional Distribution volume ratios (DVRs) of the regions of interest (ROIs) in both maternal and fetal brains using the Logan graphical analysis method.

The fitting start time (t*) was set to 30 min for fetal brain and 10 min for maternal brain based on validation of the linearity in the individual plots. DVR estimated with high fitting error (relative standard error > 10%) were excluded. To investigate the development of synaptic density in NHP brains, a linear mixed model was used to quantify the relationship between gestational ages and SV2A values: 𝑁𝐻𝑃 𝑆𝑉2𝐴 ∼ 1 + 𝐺𝐴/𝑤𝑒𝑒𝑘𝑠 + (1 | 𝑆𝑢𝑏𝑗𝑒𝑐𝑡𝐼𝐷). For these equations, the dependent variable was the SV2A value (either at the whole-brain or regional level), the independent variable was the gestational age at scan, and the random variables were the NHP ID to account for repeated measures. To aid interpretations with the human data, we first corrected NHP gestational ages based on the cross-species differences in pregnancy lengths:

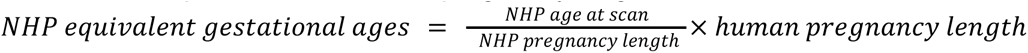

We generated the equivalent ages for each NHP scan.

### Transcriptional analysis

Gene expression microarray data were extracted from the Developmental Transcriptome Dataset (https://www.brainspan.org/). These data consist of eighteen fetal human brain samples with ages ranging from 8 to 26 weeks of gestation. Here, we focused on module C22, which is primarily related to cortical development, based on the previous results of Weighted Gene Co-expression Network Analysis by Miller et al.^12^. For each of the 11 developmental time points, we computed the mean expression value across the entire cortical mantle by averaging the participant-level normalized expression values across all available cortical samples at that age. To assess the association between transcriptional development and brain controllability, we performed Pearson’s correlation between the gene’s mean expression trajectory and the controllability values across the 11 time points. Co-developed genes were selected out with significant correlations with controllability at matched ages. Bonferroni-Holm correction for multiple comparisons was performed. Permutation tests were performed to validate the correlation between the development of gene expression and human brain controllability. Here, we permuted the temporal sequences of the gene expression 1000 times and repeated the correlation analysis with human controllability.

## Data availability

Raw Data from the Developing Human Connectome Project is publicly available at http://www.developingconnectome.org/data-release/third-data-release and can be downloaded upon request from https://biomedia.github.io/dHCP-release-notes/download.html. The raw data of the longitudinal scan of one pregnant human is publicly available at https://openneuro.org/datasets/ds005299. Gene expression data is available from (https://www.brainspan.org/).

## Code availability

Preprocessing code can be found at https://brain.labsolver.org/hcp_d2.html. Code to calculate controllability is available https://complexsystemsupenn.com/s/controllability_code-smb8.zip. Custom analysis code is available at https://github.com/huiliii/fetal_control.

## Acknowledgements

This work is supported by NIH(R21MH120615). Data were provided by the developing Human Connectome Project, KCL-Imperial-Oxford Consortium funded by the European Research Council under the European Union Seventh Framework Programme (FP/2007-2013) / ERC Grant Agreement no. [319456]. We are grateful to the families who generously supported this trial.

## Author contributions

H.S.: conceptualization, methodology, investigation, result visualization, and writing-original draft. T.T., S.M., H.L. and C.C..: investigation and result visualization. S.R., A.D., K.F., M.S. and S.G.: conceptualization and methodology; R.C.: conceptualization, writing-review and editing; D.S.: conceptualization, supervision and writing-review and editing.

## References

1. Tau, G. Z. & Peterson, B. S. Normal Development of Brain Circuits. Neuropsychopharmacol 35, 147–168 (2010).

2. Hasegawa, M. et al. Development of myelination in the human fetal and infant cerebrum: A myelin basic protein immunohistochemical study. Brain and Development 14, 1–6 (1992).

3. van den Heuvel, M. I., et al. Hubs in the human fetal brain network. Developmental Cognitive Neuroscience 30, 108–115 (2018).

4. Thomason, M. E. Development of Brain Networks In Utero: Relevance for Common Neural Disorders. Biological Psychiatry 88, 40–50 (2020).

5. Cao, M. et al. Early Development of Functional Network Segregation Revealed by Connectomic Analysis of the Preterm Human Brain. Cereb. Cortex bhw038 (2016) doi:10.1093/cercor/bhw038.

6. Gu, S. et al. Controllability of structural brain networks. Nat Commun 6, 8414 (2015).

7. Tang, E. & Bassett, D. S. *Colloquium* : Control of dynamics in brain networks. Rev. Mod. Phys. 90, 031003 (2018).

8. Cui, Z. et al. Optimization of energy state transition trajectory supports the development of executive function during youth. eLife 9, e53060 (2020).

9. Sun, H. et al. Network controllability of structural connectomes in the neonatal brain. Nat Commun 14, 5820 (2023).

10. De Asis-Cruz, J. et al. Functional brain connectivity in ex utero premature infants compared to in utero fetuses. NeuroImage 219, 117043 (2020).

11. Edwards, A. D. et al. The Developing Human Connectome Project Neonatal Data Release. Front. Neurosci. 16, (2022).

12. Miller, J. A. et al. Transcriptional landscape of the prenatal human brain. Nature 508, 199–206 (2014).

13. Rossano, S. et al. Imaging the fetal nonhuman primate brain with SV2A positron emission tomography (PET). Eur J Nucl Med Mol Imaging 49, 3679–3691 (2022).

14. Pritschet, L. et al. Neuroanatomical changes observed over the course of a human pregnancy. Nat Neurosci 27, 2253–2260 (2024).

15. Shi, F. et al. Infant Brain Atlases from Neonates to 1- and 2-Year-Olds. PLOS ONE 6, e18746 (2011).

16. Thomas Yeo, B. T., et al. The organization of the human cerebral cortex estimated by intrinsic functional connectivity. J Neurophysiol 106, 1125–1165 (2011).

17. Finnema, S. J. et al. Imaging synaptic density in the living human brain. Sci. Transl. Med. 8, (2016).

18. Hoekzema, E. et al. Pregnancy leads to long-lasting changes in human brain structure. Nat Neurosci 20, 287–296 (2017).

19. Tang, E. et al. Developmental increases in white matter network controllability support a growing diversity of brain dynamics. Nat Commun 8, 1252 (2017).

20. Spear, L. P. Adolescent Neurodevelopment. Journal of Adolescent Health 52, S7–S13 (2013).

21. Soman, S. K. et al. Cleaved PINK1 induces neuronal plasticity through PKA-mediated BDNF functional regulation. https://doi.org/10.1002/jnr.24854 doi:10.1002/jnr.24854.

22. Colantuoni, C. et al. Temporal dynamics and genetic control of transcription in the human prefrontal cortex. Nature 478, 519–523 (2011).

23. Ream, M. A. & Lehwald, L. Neurologic Consequences of Preterm Birth. Curr Neurol Neurosci Rep 18, 48 (2018).

24. Damera, S. R., et al. Regional homogeneity as a marker of sensory cortex dysmaturity in preterm infants. iScience 27, (2024).

25. Krishnan, M. L. et al. Relationship Between White Matter Apparent Diffusion Coefficients in Preterm Infants at Term-Equivalent Age and Developmental Outcome at 2 Years. Pediatrics 120, e604–e609 (2007).

26. Ball, G. et al. Development of cortical microstructure in the preterm human brain. Proceedings of the National Academy of Sciences 110, 9541–9546 (2013).

27. Bouyssi-Kobar, M. et al. Regional microstructural organization of the cerebral cortex is affected by preterm birth. NeuroImage: Clinical 18, 871–880 (2018).

28. Wilson, S. et al. Development of human white matter pathways in utero over the second and third trimester. Proceedings of the National Academy of Sciences 118, e2023598118 (2021).

29. Kisilevsky, B. S. & Hains, S. M. J. Onset and maturation of fetal heart rate response to the mother’s voice over late gestation. Developmental Science 14, 214–223 (2011).

30. Meredith Weiss, S., Aydin, E., Lloyd-Fox, S. & Johnson, M. H. Trajectories of brain and behaviour development in the womb, at birth and through infancy. Nat Hum Behav https://doi.org/10.1038/s41562-024-01896-7 (2024) doi:10.1038/s41562-024-01896-7.

31. Wass, S. V. et al. Parental neural responsivity to infants’ visual attention: How mature brains influence immature brains during social interaction. PLOS Biology 16, e2006328 (2018).

32. Karolis, V. R. et al. Maturational networks of human fetal brain activity reveal emerging connectivity patterns prior to ex-utero exposure. Commun Biol 6, 661 (2023).

33. Dufford, A. J., Spann, M. & Scheinost, D. How prenatal exposures shape the infant brain: Insights from infant neuroimaging studies. Neuroscience & Biobehavioral Reviews 131, 47–58 (2021).

34. (ISMRM 2019) The developing Human Connectome Project (dHCP): fetal acquisition protocol. https://archive.ismrm.org/2019/0244.html.

35. Omidvari, N. et al. Quantitative Accuracy Assessment of the NeuroEXPLORER for Diverse Imaging Applications: Moving Beyond Standard Evaluations. Journal of Nuclear Medicine 66, 150–157 (2025).

36. Li, H. et al. Performance Characteristics of the NeuroEXPLORER, a Next-Generation Human Brain PET/CT Imager. Journal of Nuclear Medicine 65, 1320–1326 (2024).

37. Hughes, E. J. et al. A dedicated neonatal brain imaging system. Magnetic Resonance in Medicine 78, 794–804 (2017).

38. (ISMRM 2018) Multiband zoom TSE imaging: increasing efficiency with multiband tip-back preparation pulses. https://archive.ismrm.org/2018/0062.html.

39. Kuklisova-Murgasova, M., Quaghebeur, G., Rutherford, M. A., Hajnal, J. V. & Schnabel, J. A. Reconstruction of fetal brain MRI with intensity matching and complete outlier removal. Medical Image Analysis 16, 1550–1564 (2012).

40. (ISMRM 2018) Spin And Field Echo (SAFE) dynamic field correction in 3T fetal EPI. https://archive.ismrm.org/2018/0208.html.

41. Christiaens, D. et al. Scattered slice SHARD reconstruction for motion correction in multi-shell diffusion MRI. NeuroImage 225, 117437 (2021).

42. Cordero-Grande, L., Hughes, E. J., Hutter, J., Price, A. N. & Hajnal, J. V. Three-dimensional motion corrected sensitivity encoding reconstruction for multi-shot multi-slice MRI: Application to neonatal brain imaging. Magnetic Resonance in Medicine 79, 1365–1376 (2018).

43. Veraart, J. et al. Denoising of diffusion MRI using random matrix theory. NeuroImage 142, 394–406 (2016).

44. Tustison, N. J. et al. N4ITK: Improved N3 Bias Correction. IEEE Transactions on Medical Imaging 29, 1310–1320 (2010).

45. Gholipour, A. et al. A normative spatiotemporal MRI atlas of the fetal brain for automatic segmentation and analysis of early brain growth. Sci Rep 7, 476 (2017).

46. Yeh, F.-C. et al. Differential tractography as a track-based biomarker for neuronal injury. NeuroImage 202, 116131 (2019).

47. Yeh, F.-C., Wedeen, V. J. & Tseng, W.-Y. I. Generalized $ q$-Sampling Imaging. IEEE Transactions on Medical Imaging 29, 1626–1635 (2010).

48. Towns, J. et al. XSEDE: Accelerating Scientific Discovery. Comput. Sci. Eng. 16, 62–74 (2014).

49. Jiang, H., van Zijl, P. C. M., Kim, J., Pearlson, G. D. & Mori, S. DtiStudio: Resource program for diffusion tensor computation and fiber bundle tracking. Computer Methods and Programs in Biomedicine 81, 106–116 (2006).

50. Rolls, E. T., Joliot, M. & Tzourio-Mazoyer, N. Implementation of a new parcellation of the orbitofrontal cortex in the automated anatomical labeling atlas. NeuroImage 122, 1–5 (2015).

51. Nozari, E. et al. Macroscopic resting-state brain dynamics are best described by linear models. *Nat*. Biomed. Eng 8, 68–84 (2024).

52. Gu, S. et al. Controllability of structural brain networks. Nat Commun 6, 8414 (2015).

53. Young, J. T. et al. The UNC-Wisconsin Rhesus Macaque Neurodevelopment Database: A Structural MRI and DTI Database of Early Postnatal Development. Front. Neurosci. 11, (2017).

54. Jenkinson, M., Beckmann, C. F., Behrens, T. E. J., Woolrich, M. W. & Smith, S. M. FSL. NeuroImage 62, 782–790 (2012).

55. Rossano, S. et al. Feasibility of imaging synaptic density in the human spinal cord using [11C]UCB-J PET. EJNMMI Phys 9, 32 (2022).

56. Nabulsi, N. B. et al. Synthesis and Preclinical Evaluation of 11C-UCB-J as a PET Tracer for Imaging the Synaptic Vesicle Glycoprotein 2A in the Brain. Journal of Nuclear Medicine 57, 777–784 (2016).

