## Supplementary Information for "Fetal network controllability co-develops with synaptic development and synchronizes with maternal network controllability during pregnancy" (from another version of this preprint: version 1, 2025-09-03)

Table S1. GAMLSS models for perinatal brain controllability changes in the canonical networks

| | *F* | *p* | *Rsq.(adj)* | Deviance explained |
|---|---|---|---|---|
| Whole-brain | 77.01 | <2e-16 | 0.382 | 38.60% |
| Visual | 35.78 | <2e-16 | 0.237 | 24.30% |
| Somatomotor | 10.45 | <2e-16 | 0.0892 | 9.76% |
| Dorsal Attention | 50.49 | <2e-16 | 0.336 | 34.30% |
| Ventral Attention | 4.23 | 8.02E-05 | 0.0338 | 4.26% |
| Limbic | 7.759 | <2e-16 | 0.0686 | 7.72% |
| Frontoparietal | 7.964 | <2e-16 | 0.068 | 7.67% |
| Default Mode | 10.9 | <2e-16 | 0.08 | 8.63% |
| Subcortical | 4.82 | 2.24E-04 | 0.027 | 3.15% |

Table S2. T-statistics of brain controllability between fetuses and preterm infants at the same age during the third trimester

| Gestational Weeks | N_fetal | N_preterm | *t* | *p* | *pFDR* |
|---|---|---|---|---|---|
| 29 | 20 | 7 | -4.27 | 2.49e-04 | 5.60e-04 |
| 30 | 13 | 4 | -1.57 | 0.14 | 0.14 |
| 31 | 16 | 8 | -2.19 | 0.04 | 0.045 |
| 32 | 15 | 13 | -2.41 | 0.024 | 0.032 |
| 33 | 14 | 16 | -3.07 | 0.0047 | 0.0085 |
| 34 | 7 | 17 | -2.71 | 0.013 | 0.020 |
| 35 | 7 | 36 | -5.85 | 7.04e-07 | 6.34e-06 |
| 36 | 5 | 21 | -4.62 | 1.10e-04 | 3.30e-04 |
| 37 | 3 | 9 | -6.3 | 8.87e-05 | 3.30e-04 |

Table S3. Development of NHP fetal brain sv2a values during the third trimester

| | $\beta$ | *p* | SE | 95% CI |
|---|---|---|---|---|
| Frontal | 58.58 | 0.0074 | 15.72 | [21.415,95.754] |
| Temporal | 30.74 | 0.30 | 27.40 | [-34.054,95.525] |
| Occipital | 57.12 | 0.034 | 21.74 | [5.709,108.526] |
| Parietal | 47.32 | 0.015 | 14.76 | [12.420,82.220] |

Table S4. GAMLSS models for maternal brain controllability changes in the canonical networks

| | *F* | *p* | *Rsq.(adj)* | Deviance explained |
|---|---|---|---|---|
| Whole-brain | 4.376 | 0.0167 | 0.402 | 47.30% |
| Visual | 3.474 | 0.0787 | 0.115 | 16.20% |
| Somatomotor | 5.902 | 0.00883 | 0.384 | 43.50% |
| Dorsal Attention | 3.415 | 0.0378 | 0.336 | 41.30% |
| Ventral Attention | 2.137 | 0.108 | 0.179 | 24.40% |
| Limbic | 0.74 | 0.621 | 0.0583 | 26.90% |
| Frontoparietal | 4.19 | 0.0127 | 0.527 | 63.90% |
| Default Mode | 2.661 | 0.0825 | 0.275 | 36.90% |
| Subcortical | 7.641 | 0.00173 | 0.564 | 62.60% |

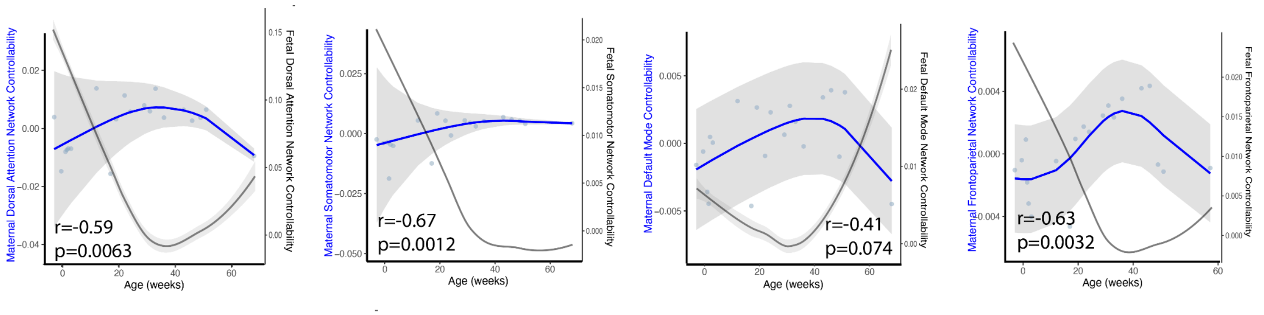


**Fig. S1.** At the network level, the controllability of the four canonical networks in the maternal brain followed an opposite trend compared to that of the fetal brain.

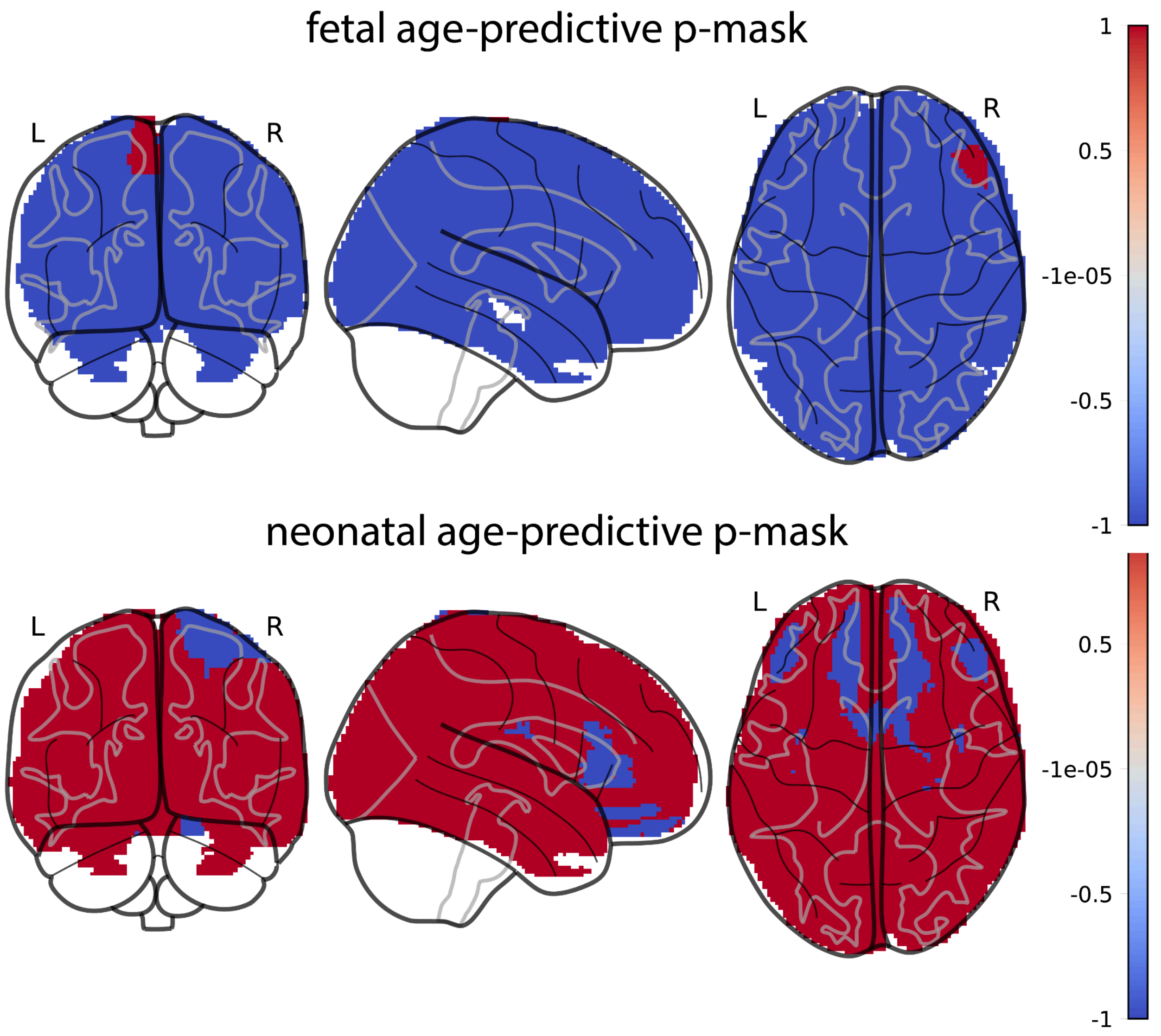


**Fig. S2 p-masks for age prediction with fetal and neonatal controllability.**
